# Workshops increase invasive species records to EDDMapS in the Mid-Atlantic United States

**DOI:** 10.1101/620138

**Authors:** Rebekah D. Wallace, Charles T. Bargeron, Jil Swearingen, Joseph H. LaForest, David J. Moorhead, Xuelin Luo

## Abstract

As the prevalence of smartphones and smartphone applications has penetrated the cellular phone market, and the capabilities of features within smartphones allowed for the replacement of commonly used mapping and identification tools, Bugwood developers began to release smartphone applications for reporting invasive species. To encourage and demonstrate reporting, workshops have been held to walk users through these apps’ features and overall EDDMapS capabilities. To ascertain the effectiveness of workshops in encouraging reporting via smartphone apps and the EDDMapS MAEDN website, a General Linear Model was used to evaluate the number of records received across the 2012-2017 time frame and evaluate if there was a difference in the number of records in the years MAEDN workshops were held at the Mid-Atlantic Invasive Plant Council conferences and workshops and years no MAEDN workshop was held. The model indicated that the workshops and time were significant factors in the number of records submitted. Being able to confidently demonstrate the effect that workshops can positively impact data recruitment can aid in funding for outreach and educational events.

## Introduction

A primary problem associated with education and outreach is the lack of ability to quantitatively assess impact. Many of these events rely upon surveys or testing for analyzing the effectiveness of the event. With certain topics, it is becoming more possible to precisely gauge if the event had an impact without relying upon a survey. When promoting invasive species mapping, it is now possible to evaluate if attendees find and map invasive species by encouraging the use of a singular application as a method of reporting. This allows tracking of invasive species mapping in response to invasive species outreach events.

The Early Detection and Distribution Mapping System (EDDMapS) is a database-driven website for invasive species occurrence reporting, data aggregation, data visualization, and source to download publicly available records and maps was released in 2005 and was initially focused on invasive plant mapping in the Southeast U.S. EDDMapS has now expanded to include mapping invasive insects, wildlife, and diseases across North America [1]. EDDMapS was developed with two primary purposes, to aggregate data from existing sources and for web-based reporting of new occurrences. The web reporting form’s fields and data formats were designed to conform to the North American Invasive Plant Mapping Standards [2], so that professionals and citizen scientists alike could directly contribute to the database. All data entered to the database is verified and, at the discretion of the verifier, publicly available for visualizations, download, reuse, and analysis. While the majority of data entered each year comes from large datasets contributed from federal and state agencies and universities, records from individuals and regional projects entered through the EDDMapS-powered websites and smartphone apps are very important for identifying new and expanding invasive species populations. The University of Georgia – Center for Invasive Species and Ecosystem Health (Bugwood) works with regional projects throughout the U.S. and continues to leverage new technology for invasive species mapping and education.

As technology advances, new developments are often evaluated for applications in all types of fields. Most recently, the cell phone has evolved into the smartphone and has become an important tool that is useful for more than just calling or texting. Inherent features on smartphones such as the camera, GPS location, internet connection, and downloadable apps, have turned the phone into an increasingly useful tool for personal and professional life. This utility has led to their overwhelming adoption among adults in the U.S. As of 2014, 64% of U.S. adults owned a smartphone and 46% said that their smartphone is something that they “couldn’t live without.” Owners feel that the phone gives them a sense of freedom (70%) rather than viewing it as a leash (30%) [3]. In fact, smartphone ownership in the 18-29 age group is at 85% and decreases with increasing age; 30-49 at 79%, 50-64 at 54% and 65+ at 27% [3]. Fully 78-80% of smartphone owners, ages 18-50+, report that their smartphones make them feel that they are productive [3]. While the native features have provided general increases in productivity, the downloadable apps have added specific value to the owners’ experience with the number and diversity of apps targeted as specific tasks increasing each year.

Before smartphone mapping technology was available for invasive plants, surveyors needed to pack a GPS unit, camera, identification guides, notebook/surveying sheets, pens, and potentially even more equipment for any specialized testing or sampling. An accuracy comparison of smartphones representative of the two operating systems and a commonly used GPS unit show that while the smartphones had a higher median location error, the Droid X with 5.2 m and the iPhone 4 with 4.6 m, they were not far from the and Garmin eTrex Legend H (2.4 m) [4]. In evaluating the quality of the smartphone camera, Hagman and Riedberg [5] compared an Apple iPhone 5 and a Samsung Galaxy S5 with a Nikon D700 in a typical medical photography situation and found the smartphones to perform well against the expensive, single function DSLR camera. As the features on the smartphones have become better, more precise, and comparable to the single-function units, surveyors have become more comfortable using these convenient tools.

The Mid-Atlantic Invasive Plant Council (MAIPC) conferences and workshops, hereafter referred to as MAIPC events, tend to draw people who are managers of lands of various size and ownership, program coordinators, volunteer managers, and others involved in the invasive species communities as professionals or individuals. The aim of the EDDMapS training sessions at the MAIPC events have been to show how to use the websites and the smartphone apps for reporting, as well as how to navigate around the websites for data querying, accessing maps, aiding in identification, and other tools and features. For those in charge of large areas or managing a crew, they can use the tools from EDDMapS to aid in their invasive species monitoring and control efforts. The attendees of these workshops are learning to become proficient users of EDDMapS and can take what they learn back to their program and train their crew or volunteers, in a “train-the-trainer” or “pyramidal” style training scheme [6]. These methods have already proven effective in other programs, such as training people who oversee others in a healthcare setting to learn how to interact with people with behavioral issues and in establishing proper procedures (such as cardiopulmonary resuscitation and other techniques) [7–8]. This model is also more cost and time efficient for the primary trainer (in these workshops, EDDMapS’ staff) to train the land managers, and the land managers to train their crew members, than for individual trainings to take place at multiple locations, and it has been proven in other studies that there is relatively high information retention rate for this training method [9–10]. However, in the training, the attendees are also assessing if they wish to use EDDMapS in their programs.

Many factors can influence if an individual participates in a crowd sourcing scientific data collection/categorization program, but often these studies’ parameters are citizen scientists in a virtual or digital space that may have a gamification or reward system, such as with the Galaxy Zoo and FoldIt programs. A citizen scientist’s motivations for participating in these programs have been largely found to be desire to contribute to scientific research and interest in science [11–12]. Studies on volunteerism in environmental conservation efforts found that learning and contribution to conservation were listed as strong motivations, with career-related reasons much weaker [13–14]. However, these studies did not evaluate a professional’s motivations when using these programs, especially when participation would potentially benefit them professionally. The MAIPC events allowed the attendees, many of which are professionals, to experience the MAEDN website and apps with their features fully explained so that they could evaluate their usefulness.

With some regions, advertisement to the public was a priority to encourage invasive species reporting. The first EDDMapS developed reporting application was IveGot1, created in conjunction with Everglades Cooperative Invasive Species Management Area (Everglades CISMA) with funding by National Park Service – Everglades National Park [15]. The purpose of the app was for public awareness as much as it was for encouraging reporting of invasive species, primarily pythons and other reptiles [15]. The IveGot1 campaign has been deemed a success, as determined by increasing numbers of records from the public, due to support by local CISMAs and other programs across the state, extension publications, and marketing via billboards, print advertisements, bumper stickers and more [15]. A similar tactic was used by the Missouri River Watershed Coalition/EDDMapS West region for increasing awareness and encouraging reporting of invasive species. That region’s funding came from a variety of sources and was used in part to create advertisements and educational materials, such as print news articles, interviews, guides, and videos on invasive species [16–18].

## Methods

Whereas the EDDMapS West/MRWC apps likely benefitted from all of the advertisement about invasive species, this method of outreach was not replicated in all regions. In the EDDMapS Mid-Atlantic Early Detection Network (MAEDN) region, the primary method of engaging the public in the regional EDDMapS reporting program was to advertise for and conduct workshops. The MAEDN region encompasses Delaware, Maryland, New Jersey, New York, Pennsylvania, Virginia, West Virginia and the District of Columbia (Fig 1). The MAIPC events were held across the region in every state and were the main outreach events and method for invasive species awareness, education, and reporter training. MAPIC events were focused on reaching the specific communities already interested in invasive species, by advertising the event through invasive species email mailing lists and direct contact with individual groups involved in invasive species.

**Figure 1.**
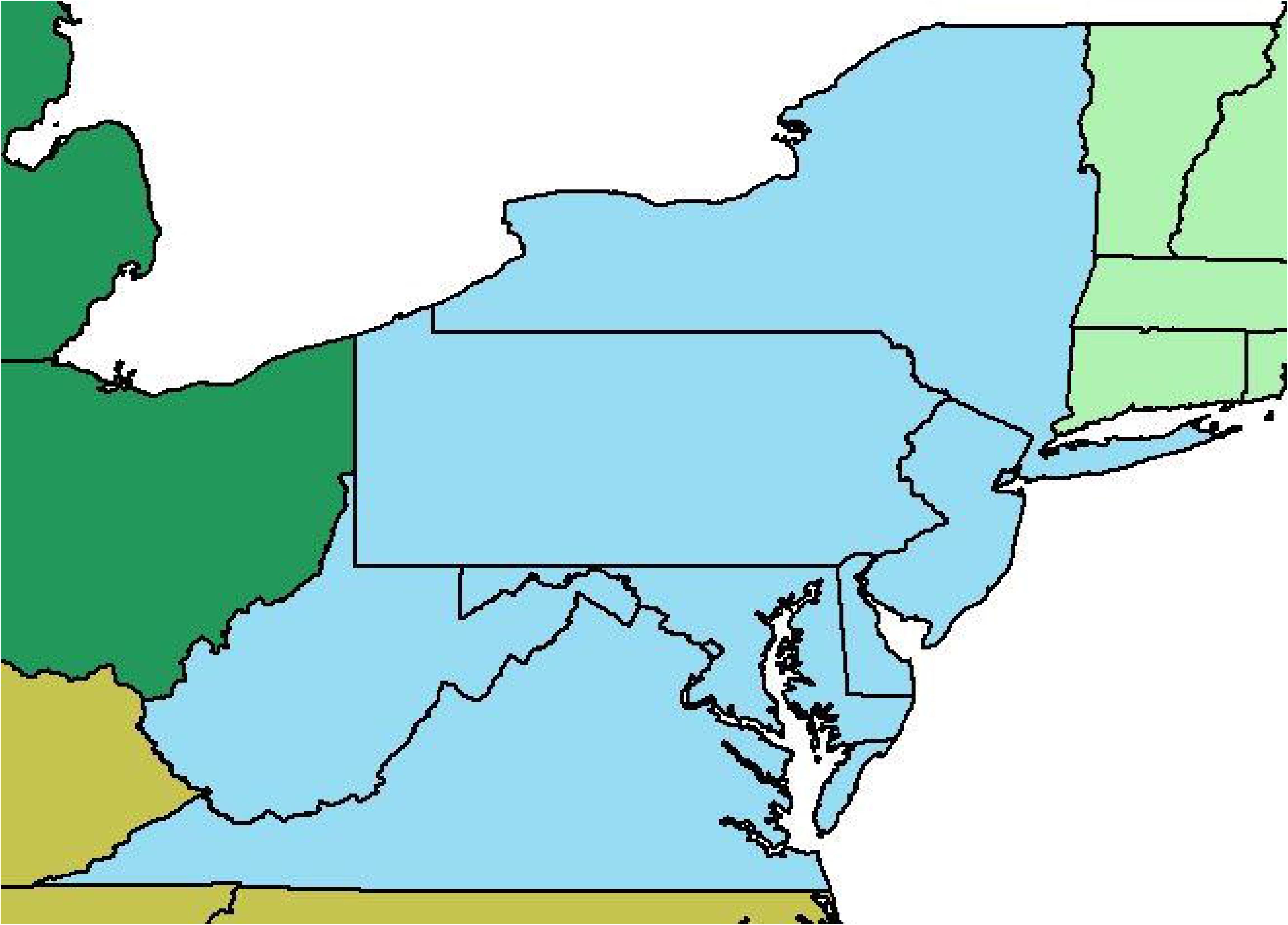
Mid-Atlantic Early Detection Network Region. Region of the United States that is within the Mid-Atlantic Early Detection Network is colored blue and includes Delaware, Maryland, New Jersey, New York, Pennsylvania, Virginia, West Virginia, and the District of Columbia.

The MAEDN website and smartphone applications (iOS and Android platforms) were originally launched in April and May 2012, respectively, with updates and new versions being released as time has passed. Invasive species cooperatives, parks, forests, wildlife refuges, and other groups focused on natural areas within the MAEDN region have been concerned with invasive species for many years. However, reporting of invasive species was cumbersome until the phone app became available, the data was not broadly aggregated, and there was no singular, easy-to-use reporting tool. The MAEDN app and websites were built to fulfill this need.

### MAIPC Events

Since the regional website and apps were launched, Bugwood Staff and MAEDN project partners have presented at several of the MAIPC events and conducted demonstrations on the features and, specifically, on reporting using the apps and website. The MAIPC events were held between July and September each year from 2012-2017, with MAEDN workshops conducted on September 13, 2012 in Washington DC, on August 1, 2013 in Shepherdstown, WV, on July 30, 2014 in Washington DC, and on July 11, 2016 in Laurel, MD. EDDMapS’ staff presented at the MAIPC events in 2012, 2014, and 2016 and a project partner presented in 2013.

The MAIPC events had presentation sessions entirely revolving around identification, reporting, control, mapping, emerging invasive species and more, which EDDMapS’ staff was involved in soliciting and curating topics of presentations during several of the years. For the MAEDN workshops that were conducted at MAIPC events, presenters were allotted 30-60 minutes to demonstrate the EDDMapS MAEDN website and apps, and cover existing and new features added each year as technology evolved. Presentations were a combination of a prepared projected slide presentation and live demonstration of the applications and website. Each workshop also included a separate hands-on field exercise in a nearby area to allow attendees to practice navigating through the apps and reporting invasive species using the app. Presenters would roam the area answering questions and offer attendees guidance as needed during the field exercise. After the field exercise, attendees were regrouped with the presenter to discuss their experience, assess mapping results, answer questions, address any issues that arose, and discuss the review process and publishing of records after the data are sent to the EDDMapS database. These exercises helped to train the attendees on the MAEDN apps, better understand the various aspects of the MAEDN apps and the EDDMapS website, and provided them the knowledge they needed for their invasive species management programs. Attendees also gained the skills needed to teach their crews, staff, and volunteers how to incorporate the tools into their existing programs.

### MAEDN Reporters

Similar, to other species occurrence reporting applications (e.g., iNaturalist, eBird, etc.) there are no educational or experiential criteria that must be met to participate and report species. One simply needs to register for an account to submit records and users are encouraged to participate to increase the volume of occurrence data [19–20]. Reporters are using the MAEDN app to document invasive species occurrences and are informed through training and online documentation that data, once checked and verified to be correct, will be uploaded to the database and available as maps and for download. As with all of EDDMapS-based citizen science reporting apps, the MAEDN app is free to use; anyone can register for an account at any point to use the app to report invasive species. The primary method for increasing the number of reporters in the MAEDN region was through the MAIPC events. While MAIPC events are open to anyone interested, the advertising for them was mainly done through email mailing lists from MAIPC, state invasive species councils, state and local agencies, federal agencies, University Extension Services, and the MAIPC and other websites. As a result, the events are most often attended by those already interested or invested in the invasive species community either as volunteers, researchers, or professionals for state and federal agencies, private companies, non-profits, academia, public lands, and more [21]. Attending these events and electing to become an EDDMapS reporter is a self-selecting process. Not all who attend the MAEDN workshops will create accounts or contribute records. Professionals and other programs can have very defined needs in their tools and usability and compatibility will drive them to use the system which will best work with their needs and to reject systems missing key components. Some considerations for a tool useful for professionals documenting invasive species occurrences include fit into their existing program, database compatibility, documentation standards, etc.

MAEDN and other EDDMapS’ regional apps are used by professionals as well as citizen scientists, with citizen scientists primarily using them during recreational or volunteer activities and professionals using them to document invasive species occurrences as part of their professional activities, for example, to develop management plans. Registration for an EDDMapS account is very simple, requiring a first name, last name, email address, and user-created password. Other information can be added either through the registration process or by the user editing their EDDMapS profile (i.e., organization, unit/department, phone number, country, state, inclusion within a specific user group, self-assessed knowledge level for various species categories, if they have completed training in the First Detector program, etc.). Only first name, last name, and organization are included in the publicly available and verified invasive species record.

### Submitting A Record

The records enter the EDDMapS database via websites, web embedded forms, reporting API, or through data files sent to the EDDMapS Data Coordinator and are uploaded as Not Reviewed and Not Public. At minimum, a record must have a Reporter, Species Name, Location, and Observation Date. Reporter name is the name of the person logged into the EDDMapS account when the report is created or the name provided from those submitting bulk data to the EDDMapS Data Coordinator. Species name is selected from a drop-down list that is embedded within the reporting app or the EDDMapS website. Observation date is defaulted in the apps and websites to the current data, but that is able to be edited. Location must at least be a state/province and a county/county-equivalent (e.g., parish, borough, municipality, independent cities, etc.), though GPS coordinates are preferred as a point, line, multiline, polygon, or multipolygon. Additional fields to provide information on extent, abundance, species/population characteristics, ecosystem, and more are available on the EDDMapS website and a shorter form is available through the MAEDN app. Both the website and apps encourage including images of the species to be added to the report.

Each Bugwood-developed citizen science reporting app and website (e.g., MAEDN, IveGot1, EDDMapS West, etc.) upload data in a state of Not Reviewed and Not Public when it enters the database. The record is verified by experts in that subject and are often located within that geographic region (e.g., county, state, CISMA/CWMA region, etc.).

### EDDMapS Record Review Process

The only data included for evaluation were those that were at the status of Reviewed and Publicly Available. The names of the EDDMapS record verifiers and the review process are both publicly available, verifiers can be found on the Report Verifier Lookup tool on the EDDMapS website (https://www.eddmaps.org/tools/verifierlookup/) and the verification process is published on the Bugwood Presents website [22] and a link to that presentation is also available on the EDDMapS Tools & Training page.

Species that are regulated by state or federal law go to the appropriate state and federal agency contacts, as there are laws regarding species classified as such. Species that are not regulated are reviewed by individuals that have education and/or extensive training in invasive species identification and these individuals are often limited to a geographic area to aid in identification credibility. Notification of new reports is sent to individuals including academic researchers, members of invasive species programs (e.g., CISMAs, CWMAs, etc.), state and federal botanists and biologists whose parameters match reports (i.e., species, geographic area, project, etc.).

The verifier evaluates the data provided by the reporter in the record, soliciting additional information and/or images as needed, and submits their decision. The verifier can approve the record, change the species (if name selected by the reporter was incorrect), change the record from a positive (species detected) to a negative (species not detected) status if the species was incorrect, or reject the record. Records that are approved, species corrected, and negative are able to also have their status set to “publicly available” which allows the records to show up on maps, be available for download, and are otherwise shareable.

### Data Queried

The data in the database is constantly evolving, as data is reviewed by verifiers and/or existing records are contested/flagged the numbers can fluctuate. However, the further from the target date range, the records become more stable as there are fewer records left to review and fewer reviewed records are subject to contesting and flagging. All data for EDDMapS are housed on off-site servers with Amazon Web Services for data preservation and security. Most of the information associated directly with a record is publicly available upon verification, however, the only data available with the record about the reporter is their name and organization (if they chose to enter organization) for display on the website and in downloaded data files.

The data used in this evaluation were only those that were Reviewed and Publicly Available, and were entered via the MAEDN website or MAEDN smartphone applications with observation dates between January1, 2012 and December 31, 2017. The data were queried from the EDDMapS database on January 16, 2018 and the number of records by observation date per month were saved in a Microsoft Excel workbook for analysis. January was deemed a safe time to query the data, as very little data is posted annually after October, even in high volume years.

The only data fields returned from the database for this analysis were: month of observation date, year of observation date, project, source type, reviewed, and actionable. All data in the EDDMapS database are, upon uploading to the database, marked with the project the data came from. This could be the website used, the app used, or the database the data was contributed from. Project allowed the query to be specific to the MAEDN project. Source type is used to note the technology used to send the data to the database, this analysis limited the sources to iOS, Android, and Website. Reviewed governs if the records were verified and actionable is the public/not public status of the record. As only data that is accurate is retained in the database, Rejected records were not included in this query.

To assess the impact of events on individual records submitted to EDDMapS in the MAEDN region, we evaluated the number of records received during June, July, August, and September (summer) of 2012-2017. Data was evaluated in SAS with a General Linear Model (GLM) expressed as:

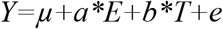

where *μ* is the intercept, *a* is decreased counts when there is no event, 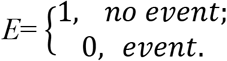, *b* is the slope of time, and *T* is time, e is random error. The number of records received during the summer of each year is the dependent variable (Records = *Y*) and the workshops is the independent variable (Event = *a*). The fitted model is *Ŷ* = −50.8229870 – 266.5790140\**E*+ 9.6626487 * Time.

## Results

The focus of the MAEDN workshops was presentation and demonstration of EDDMapS, the MAEDN website, and the MAEDN smartphone app and applicability to existing invasive species programs. The percentage of records submitted via the MAEDN smartphone app rapidly grew over the 2012-2017 timeframe. By the second year that the apps were available, they were the source of over 63% of individual records submitted and each year since has been the source of 53-86% of individual records (Table 1). This trend is also being observed in the EDDMapS database for other regions of the U.S., as both the public and professionals are finding convenience in using smartphones to report invasive species.

**Table 1.** Number of individual records submitted via MAEDN website and smartphone apps

| Year | Smartphone Records | Web Records | Total | Smartphone % | Web % |
| --- | --- | --- | --- | --- | --- |
| 2012 | 113 | 165 | 278 | 40.64748201 | 59.35251799 |
| 2013 | 683 | 386 | 1069 | 63.89148737 | 36.10851263 |
| 2014 | 645 | 440 | 1085 | 59.44700461 | 40.55299539 |
| 2015 | 500 | 431 | 931 | 53.7056928 | 46.2943072 |
| 2016 | 2525 | 389 | 2914 | 86.65065202 | 13.34934798 |
| 2017 | 1538 | 421 | 1959 | 78.50944359 | 21.49055641 |

Number of records submitted to the EDDMapS database via the MAEDN smartphone applications, the MAEDN website, the total of the apps and website, the percentage of total records coming from smartphone apps, and the percentage of total coming from the MAEDN website. Submissions via smartphone apps accounted for a higher percentage of the total individual records contributed compared to web records by the second year (2013).

The GLM fitted model has an R^2^ of .758404 and the p-value indicated that Time and Event were both significant in number of records submitted (Table 2).

**Table 2.** Type III Sum of Squares

| Source | Degrees of Freedom | F value | Pr>F |
| --- | --- | --- | --- |
| Time | 1 | 65.90 | <.0001 |
| Event | 1 | 26.46 | <.0001 |

Type III Sum of Squares shows that Time and Event are both significant factors in the number of records received from the MAEDN project area with p-value of <.0001.

Figure 2 indicates that records spike during summers in general, but the model indicates that events result in a significant increase in the number of records submitted through the MAEDN website and apps. The model also indicated that Time is a significant variable, and as groups and individuals have attended the MAEDN workshops and discovered the reporting apps and website through other means (e.g., word of mouth, media, conferences, etc.) reporting has increased over the years. This increase in records over the course of time is reflected in the linear regression line: *y*=*ax* + *b*, where *y* is the number of records, *a* is the slope of the line, *x* is Time, and *b* is the intercept.

**Figure 2.**
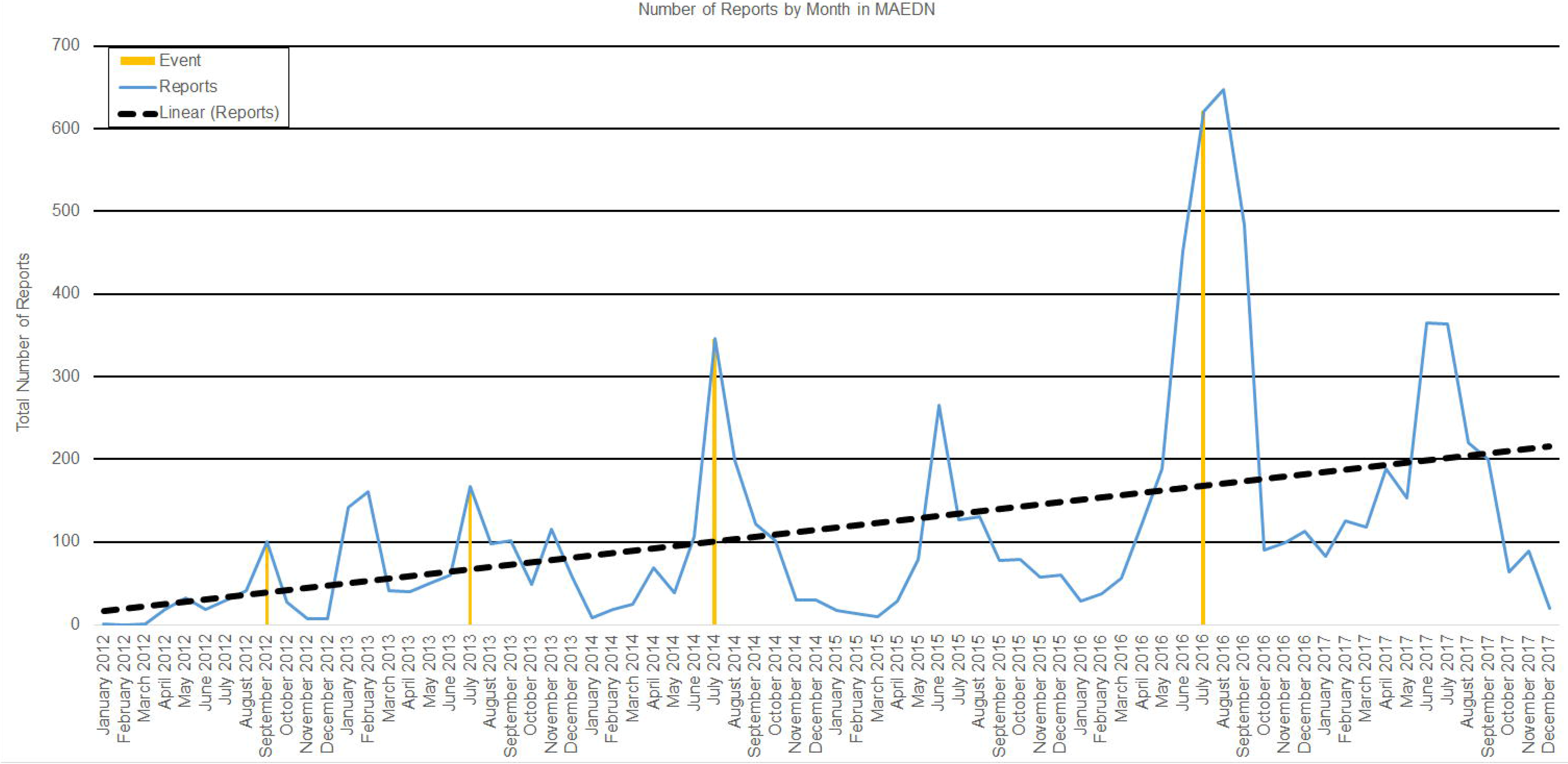
Number of Individual Records by Month in MAEDN. Caption: Number of records submitted through the MAEDN website and the MAEDN smartphone apps on a monthly basis are represented by the solid blue line. The MAEDN workshops are represented by a vertical orange line marking the month of the event. A linear regression (*y*=*ax* + *b*), where *y* is the number of records, *a* is the slope of the line, *x* is Time, and *b* is the intercept, over the analysis period was included to show increasing records over the time frame is represented by the black broken line.

A least square means (LSM) of the fitted model of the GLM was also evaluated to predict if there would have been a significant effect of a workshop on the number of records submitted in 2015 and 2017, the two years that did not have MAEDN workshops during the time frame. June was selected as this is the naturally occurring height of reporting during years that events were not held. The LSM evaluated the effect of a workshop conducted in June in 2015 and June 2017 and found that both years would have had a significant increase in records at 95% confidence limits (Table 3). This indicates that it would be beneficial to attempt to regularly host MAEDN workshops at the MAIPC events to increase awareness, and thus reporting.

**Table 3.**
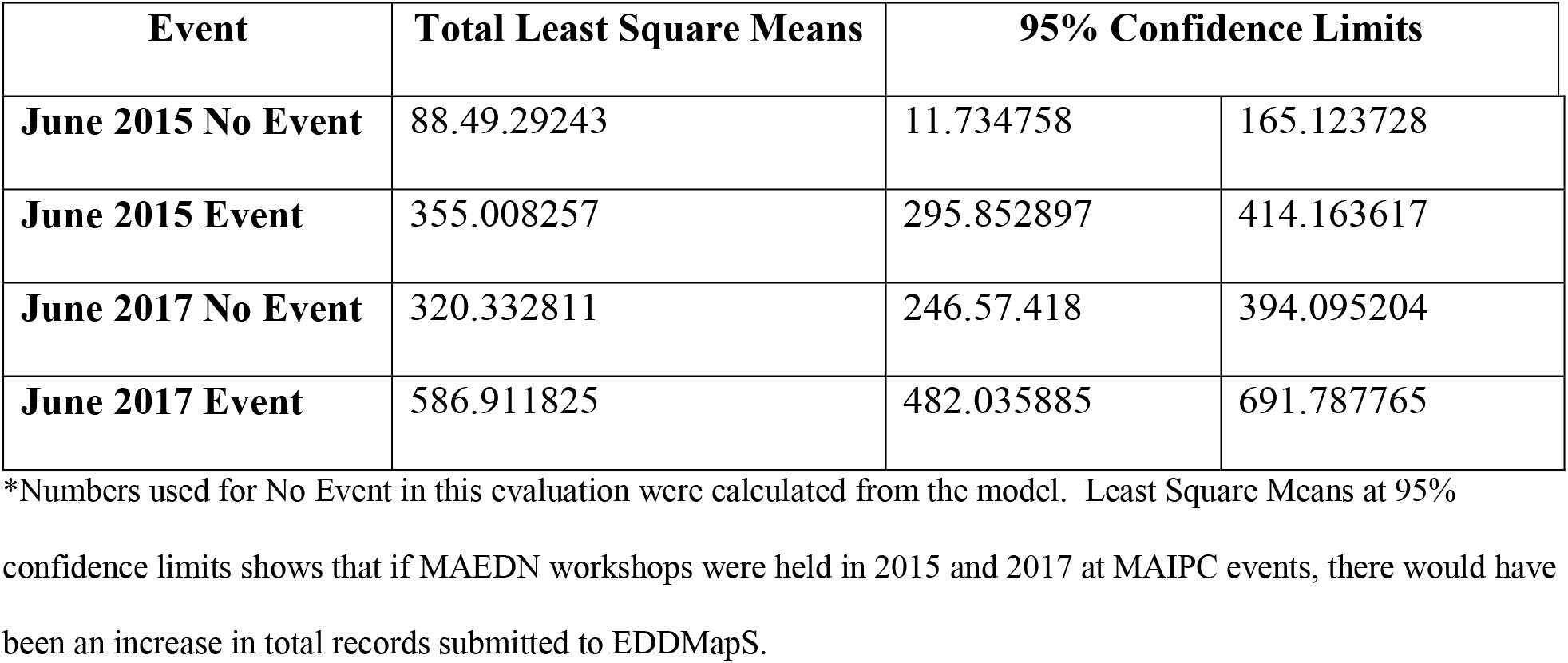
Least Square Means at June in 2015 and 2017*.

## Conclusions

A GLM analysis of records from smartphone applications and web records in the MAEDN area from 2012-2017 were evaluated to determine the effect of MAEDN workshops conducted at the MAIPC events on the number of reviewed records submitted to the EDDMapS database. Workshops on MAEDN, and specifically on the MAEDN smartphone applications, were conducted to an audience mainly comprised of landowners, professionals, and those responsible for crew, staff, or volunteers. Due to staff or partner unavailability, MAEDN workshops were not held in 2015 and 2017. The GLM indicated that Year and Event were significant in the number of records submitted through the MAEDN website and MAEDN smartphone apps. This indicates that while records are increasing each year, potentially as word spreads, through organic web searches, or by attending non-MAIPC EDDMapS presentations, the MAIPC events are also important in encouraging records in this region. The LSM indicates with a 95% confidence limit that, were workshops conducted in 2015 and 2017, similar increases in reporting would have been observed. With this information, funding and support may be able to be sought and directed to attending and demonstrating at comparable events in other regions to attempt to replicate this effect.

